# The discovery of genome-wide mutational dependence in naturally evolving populations

**DOI:** 10.1101/2022.06.24.497504

**Authors:** Anna G. Green, Roger Vargas, Maximillian G. Marin, Luca Freschi, Jiaqi Xie, Maha R. Farhat

## Abstract

**Background:** Evolutionary pressures on bacterial pathogens can result in phenotypic change including increased virulence, drug resistance, and transmissibility. Understanding the evolution of these phenotypes in nature and the multiple genetic changes needed has historically been difficult due to sparse and contemporaneous sampling. A complete picture of the evolutionary routes frequently travelled by pathogens would allow us to better understand bacterial biology and potentially forecast pathogen population shifts.

**Methods:** In this work, we develop a phylogeny-based method to assess evolutionary dependency between mutations. We apply our method to a dataset of 31,428 *Mycobacterium tuberculosis* complex (MTBC) genomes, a globally prevalent bacterial pathogen with increasing levels of antibiotic resistance.

**Results:** We find evolutionary dependency within simultaneously- and sequentially-acquired variation, and identify that genes with dependent sites are enriched in antibiotic resistance and antigenic function. We discover 20 mutations that potentiate the development of antibiotic resistance and 1,003 dependencies that evolve as a consequence antibiotic resistance. Varying by antibiotic, between 9% and 80% of resistant strains harbor a dependent mutation acquired after a resistance-conferring variant. We demonstrate that mutational dependence can not only improve prediction of phenotype (e.g. antibiotic resistance), but can also detect sequential environmental pressures on the pathogen (e.g. the pressures imposed by sequential antibiotic exposure during the course of standard multi-antibiotic treatment). Taken together, our results demonstrate the feasibility and utility of detecting dependent events in the evolution of natural populations.

Data and code available at: https://github.com/farhat-lab/DependentMutations

## Introduction

Genomic evolution of pathogenic bacteria is rapid, pervasive, and poses a serious threat to global health. The evolutionary pressure imposed by human infection creates pathogens that are more transmissible, more virulent, or more difficult to treat due to antibiotic resistance. While often attributed to single mutational events, antibiotic resistance is more complex, and high-level resistance can manifest through multiple mutations in a sequential and dependent manner. Dependency, here defined as when an initial mutation changes the likelihood of a specific subsequent mutation, may arise due to the fitness cost of initial resistance acquisition, or the action of antibiotics on multiple cellular processes (*1*, *2*). A complete understanding of the multiple, dependent mutations associated with any pathogen phenotype, including resistance, would allow us to better understand pathogen biology and potentially forecast evolution.

Traditionally the study of mutational dependence in microbial populations has relied on *in vitro* evolution experiments where populations are longitudinally sampled to determine mutational trajectories (*3*–*6*). This heavily restricts the context and breadth of evolutionary landscapes we can study. Further, resistance acquisition *in vitro* may not necessarily reflect resistance acquisition *in vivo* within a host environment. New approaches are needed to understand evolution of natural populations that will necessarily be sampled contemporaneously and be the most relevant to real-world scenarios and human health.

*Mycobacterium tuberculosis* complex (MTBC), the causative agent of tuberculosis, which displays increasing antibiotic resistance globally, is an important case study for identifying mutational dependency (*7*, *8*). While prior reports have characterized individual cases of dependent evolutionary trajectories in MTBC antibiotic resistance (*4*, *9*, *10*), a genome-wide method to detect dependent mutations generalizable to any phenotype is needed. In other bacterial species, recent work has used Potts models and regression with interaction terms to detect dependent evolution in natural populations (*11*–*13*). However, the strong linkage effects and low diversity of many pathogens, including *M. tuberculosis*, require an alternative approach **(Supplement)**. A well-suited solution to clonally evolving populations is to focus on mutations that evolve in a parallel manner across the phylogeny. This approach has been successful in detecting individual genetic effects on phenotype because it readily controls for population structure, biased sampling, and linkage across the clonal genome (*14*, *15*). Such an approach has not to date been adapted to study dependencies between pairs of mutations.

Here, we study pairs of dependent, parallelly evolving (homoplastic) mutations arising during the evolution of a natural population. We determine which mutations are more likely to occur in certain genetic backgrounds, controlling for increased uncertainty when mutations are rare. We applied our method to a dataset of 31,435 MTBC genomes spanning six major global lineages, finding that antibiotic resistance and antigen evolution are enriched among dependent mutation pairs. We observe 20 mutations that appear to potentiate the evolution of resistance to multiple different antibiotics. We quantify the number of strains in our dataset with evidence of dependent evolution occurring as a consequence of initial resistance evolution to 12 antibiotics – with lower bounds ranging from 80% for streptomycin to 9% for fluoroquinolones. We chart common manifestations of these consequential mutations after antibiotic resistance evolution, finding compensatory variation mediated through both physical interactions and metabolic pathways, multistep evolution of high-level resistance phenotypes **(Figure 1)**. Overall, our results demonstrate the promise of detecting dependent mutational events in naturally evolving pathogen populations and explore mechanistic explanations for dependencies.

**Figure 1:**
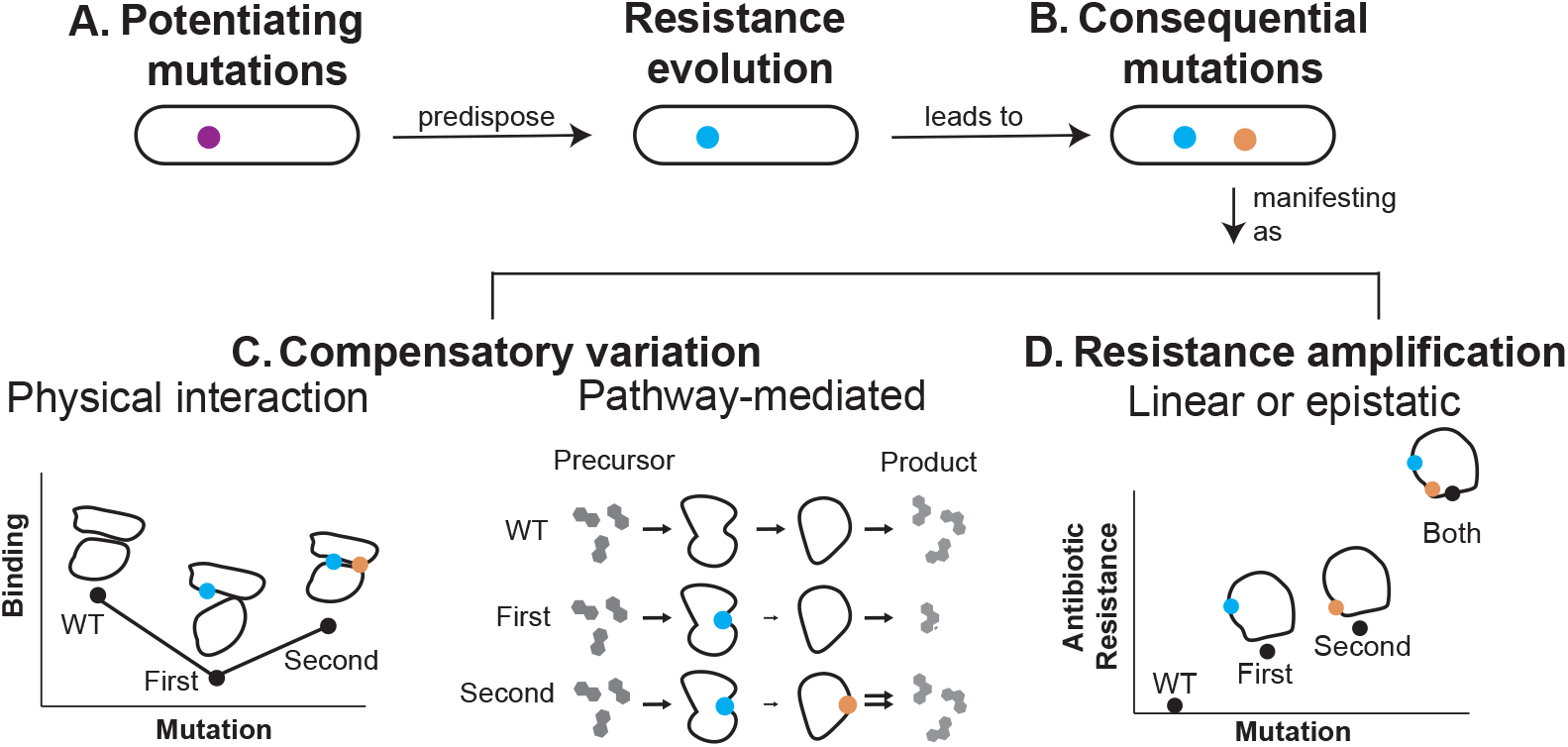
Patterns leading to detected evolutionary dependency. A simplistic framework classifying observed types of evolutionary dependencies in antibiotic resistance development. Dependencies can potentiate resistance development (A). Potentiating mutations may amplify resistance, i.e. directly influence the inhibitory concentration of the drug, or they may instead have a general effect on growth, virulence, or metabolism that increase the probability of acquisition of directly causal drug resistance mutations. After initial resistance evolution, consequential mutations (i.e, arising as a consequence of resistance) are observed and manifest through multiple mechanisms. Consequential mutations may restore fitness lost with the acquisition of resistance variants. The latter can be mediated through direct physical interactions (B) or pathway-mediated changes in related genes (C). Lastly, consequential mutations can causally amplify resistance, either through individual effects or epistatic effects such that the combination of the two variant effects is different than the sum of the individual effects (D).

## Results

### Evolutionary events in *M. tuberculosis*

We estimated the evolutionary history of 31,435 diverse MTBC strains using maximum likelihood phylogeny and ancestral sequence reconstruction, with 2,815, 8,090, 3,398, 16,931, 98, and 96 strains belonging to Lineages 1-6, respectively as recently described (*16*). Restricting our analysis here to single nucleotide polymorphisms, we observe 4,776 sites in the genome to have evolved away from the pan-susceptible ancestral state (*17*) at least five times independently **(Supplementary Data 1)**. Of these 4,776 sites, 19% are intergenic, and the remaining mutations are found in a total of 1,479 different genes. The mutations are well-distributed phylogenetically, arising in a median of three major lineages. Most mutations are relatively recent, with each mutation inherited on a median of 2.4 descendant branches.

We then categorize the homoplastic mutations in terms of their putative function: labelling mutations as antibiotic resistance associated based on a catalog of known and potential variants (*18*) and antigenic based on their presence in proteins with known epitopes (*19*, *20*) **(Methods)**. Antibiotic-associated mutations are overrepresented in our dataset of homoplastic mutations, with 5% and 17% of mutations annotated as known or possibly resistance-conferring, respectively (Chi-squared p-value < 10^-307^ for both) **(Supplementary Data 2)**. Homoplastic mutations in epitopes and epitope-containing proteins comprise 3% and 17% of the dataset, respectively, again representing a significant enrichment (Chi-square *p*-value < 10^-68^ and < 10^-39^). We find no significant enrichment for homoplastic mutations in essential genes (Chi-squared p-value > 0.1). The overrepresentation of antibiotic resistance and antigen associated homoplastic mutations suggests positive selection for these beneficial traits.

### Detecting dependencies between mutations

We develop a method to detect dependency between pairs of homoplastic mutations. We first partition the dataset into two non-mutually exclusive groups: (1) mutation pairs that occur simultaneously on the same branch at least once (N=139,048), and (2) mutation pairs that occur sequentially on subsequent branches at least once (N=1,274,446).

To test for dependencies between sequentially occurring mutations *a* and *b*, we determine if the estimated probability of mutation *a* is higher for a genetic background containing mutation *b* compared with the root ancestral background **(Figure 2, Methods)**. We detect significant evolutionary dependency for 2.0% (N=25,947) of all sequentially occurring homoplastic mutation pairs (Benjamini-Hochberg FDR < 0.01) **(Supplementary Data 3)**.

**Figure 2:**
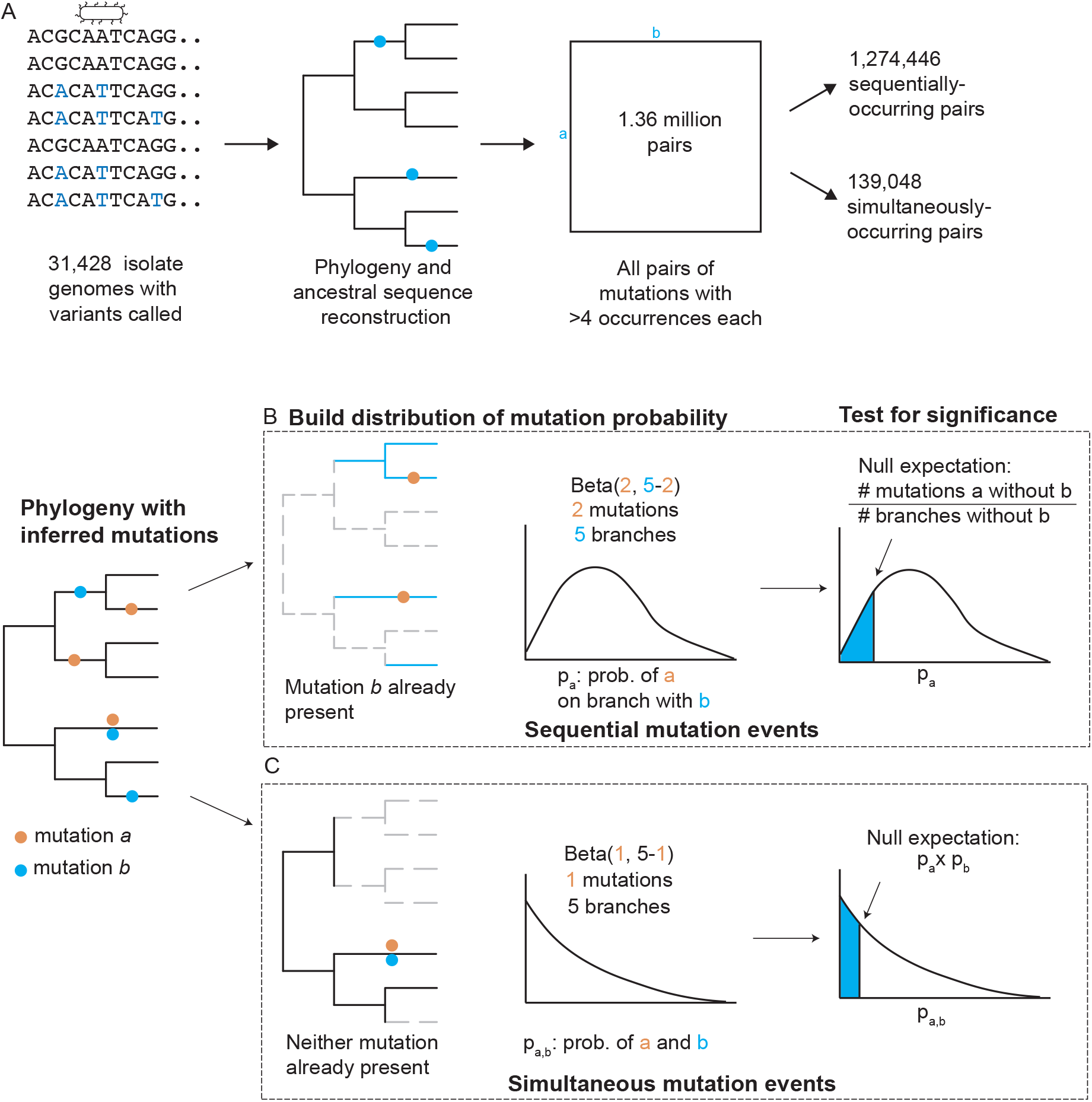
Computational workflow for finding dependencies between mutations. Workflow for finding significant dependencies between pairs of mutations. (A) We found 1,364,150 million pairs of single nucleotide polymorphisms (SNPs) across 4,776 sites that co-occur either sequential or simultaneously at least once. We began with a dataset of 31,428 isolate genomes and performed phylogeny and ancestral sequence reconstruction. We called each SNP as ancestral or derived relative to the pan-susceptible *M. tuberculosis* ancestral sequence (H37Rv), then enumerated all SNPs that arise at least 5 times independently, dividing them into pairs that appear at least once sequentially or simultaneously. (B) For sequentially occurring pairs, we determine whether the probability of mutation *a* is affected by the presence of mutation *b* by inferring the distribution of the probability of mutation *a* in the context of *b* using a Beta distribution, and then comparing it to the expected probability of mutation *a* not in the context of *b*. (C) For simultaneously occurring mutations, we determine whether the probability of observing mutations *a* and *b* simultaneously is higher than expected based on the product of the individual probabilities of mutation *a* and *b* – *ie*, assuming the two events are independent.

To test for dependencies between simultaneously occurring mutations *a* and *b*, we determine if the estimated probability of mutations *a* and *b* occurring simultaneously is higher than the estimated frequency of their co-occurrence if the two mutations were independent events **(Figure 2, Methods)**. We detect significant evolutionary dependency for 46% (N=63,271) of all simultaneously occurring homoplastic mutation pairs (Benjamini-Hochberg FDR < 0.01) **(Supplementary Data 3)**. We note the high fraction of significant pairs because simultaneous occurrence of any two mutations on a branch is unlikely.

### Sequentially occurring dependencies are enriched in antibiotic resistance function

We next annotate whether the pairs of sequentially occurring mutations are enriched in antibiotic resistance-associated or antigenic proteins. We greedily assign each pair of mutations into the following categories in the respective order: both mutations are antibiotic-associated, the first or second mutation acquired is antibiotic-associated, both mutations are antigenic, one mutation is antigenic, or other (none of the categories apply) **(Methods)**.

The pairs of sequentially occurring dependent mutations are enriched in antibiotic resistance (actual: 13.4% vs. expected: 10.2%) and antigenic categories (actual: 43.7%. vs. expected: 29.1%) compared to our expectation from the frequencies of individual SNPs (Chi-squared p-value < 10^-307^, **Supplementary Data 2**). This indicates that not only are individual antibiotic resistance and antigen-associated mutations individually under positive selection but that there are relationships between pairs of mutations that render some of them more likely to co-occur in one another’s presence. Among the top 100 hits in terms of p-value, 69% include a known resistance variant, and all of these are pairs where the known resistance-conferring mutation occurs second **(Figure 3A).**

**Figure 3:**
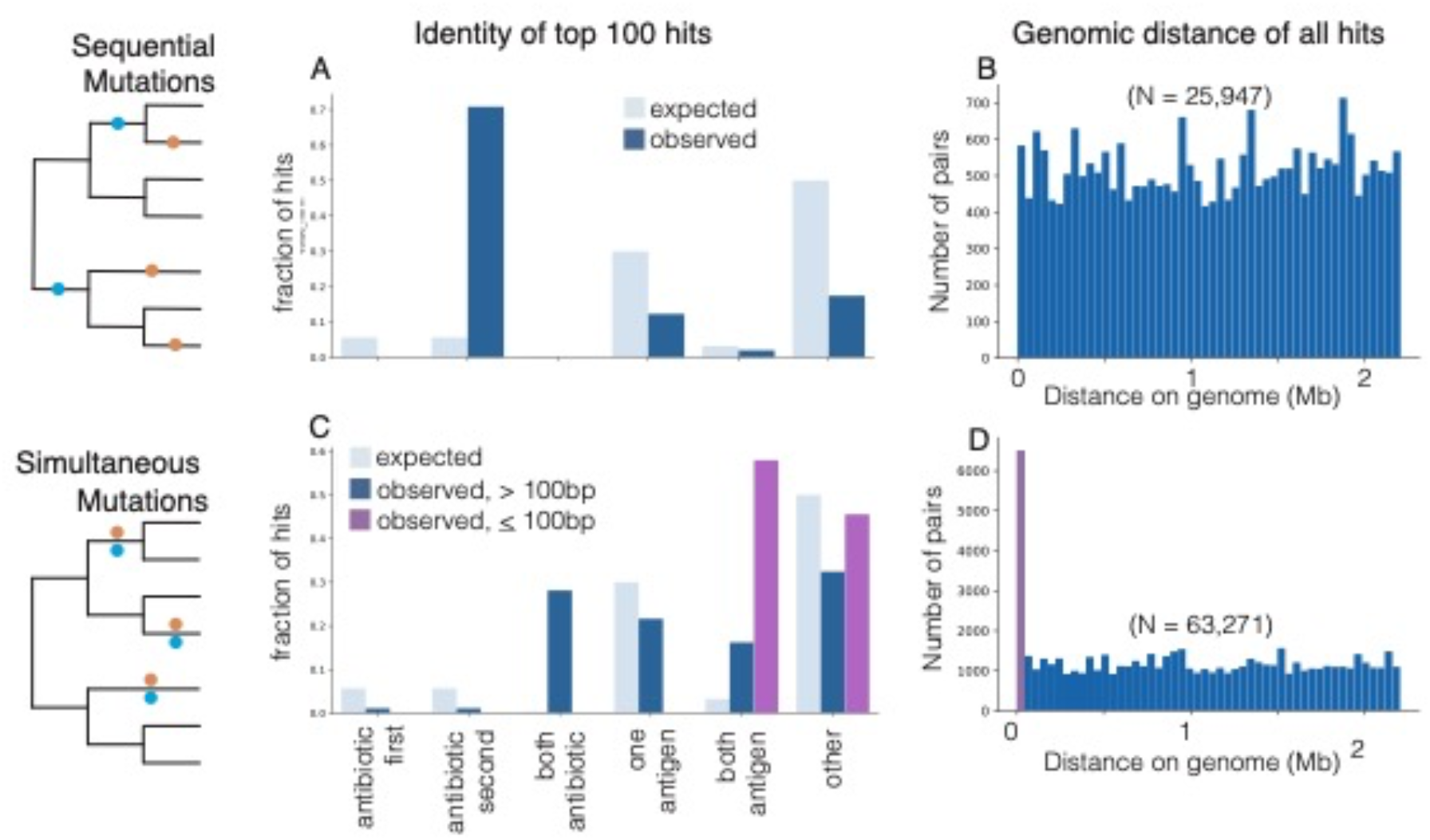
Sequential and simultaneous mutation pairs are enriched in functional categories. We determine the identity of the top 100 pairs of significant hits for (A) sequential mutation pairs and (C) simultaneous mutation pairs. We categorize mutation pairs as those where a known resistance mutation occurs, known resistance mutation occurs second, both mutations are known resistance mutations, one mutation is in a known antigen protein, both mutations are in a known antigen protein, or other category (not any of the above). For simultaneous mutations, we compute the categories for the top 100 hits found within 100 base pairs on the genome, and for the top 100 hits found outside 100 base pairs. The genomic distance in megabases of all pairs of significant dependent mutations for (B) sequential mutations and (D) simultaneous mutations are shown.

### Simultaneously occurring dependencies are enriched in antigenic function

We find that simultaneously occurring dependent pairs of homoplastic mutations are enriched in functional categories compared to our expectation from the frequencies of individual SNPs (Chi-squared p-value < 10^-307^, **Supplementary Data 2**). We then identify that simultaneously occurring dependent pairs are more likely to be in close genomic proximity than sequentially occurring dependent pairs **(Figure 3B and 3D).** Of the top 100 pairs of simultaneous dependencies, 92% are within 100 base pairs on the genome. The skewed distribution of genomic proximity suggests that these mutations are acquired simultaneously through a different mechanism than single nucleotide substitution, for example, gene conversion or recombination (*21*, *22*). Hence, we examine simultaneously occurring *proximal* mutations (<=100bp) separately from simultaneously occurring *distant* mutations (>100bp). Over 50% of the top 100 significant proximal pairs both occur in an antigenic protein **(Figure 3C)**. Among the top 100 significant distant pairs, both antigenic and antibiotic resistance-conferring pairs of mutations are overrepresented (Chi-squared p-value < 10^-89^).

### Potentiating mutations that predispose the evolution of antibiotic resistance

We examine whether particular SNPs predispose the evolution of antibiotic resistance, here called potentiator mutations, as these are of high interest for surveillance and genomic prediction. Among all 25,947 pairs of mutations with significant sequentially acquired dependency, the resistance-conferring mutation is second in 2,162. Over half of these, 1,192 of 2,162, are explained by just 20 initial mutations which we here define as potentiators because they lead to over 30 different resistance-associated mutations each **(Table 1)**, indicating that they do not predispose the strains to the evolution of resistance to a particular antibiotic, but rather predispose to resistance phenotypes in general.

**Table 1:**
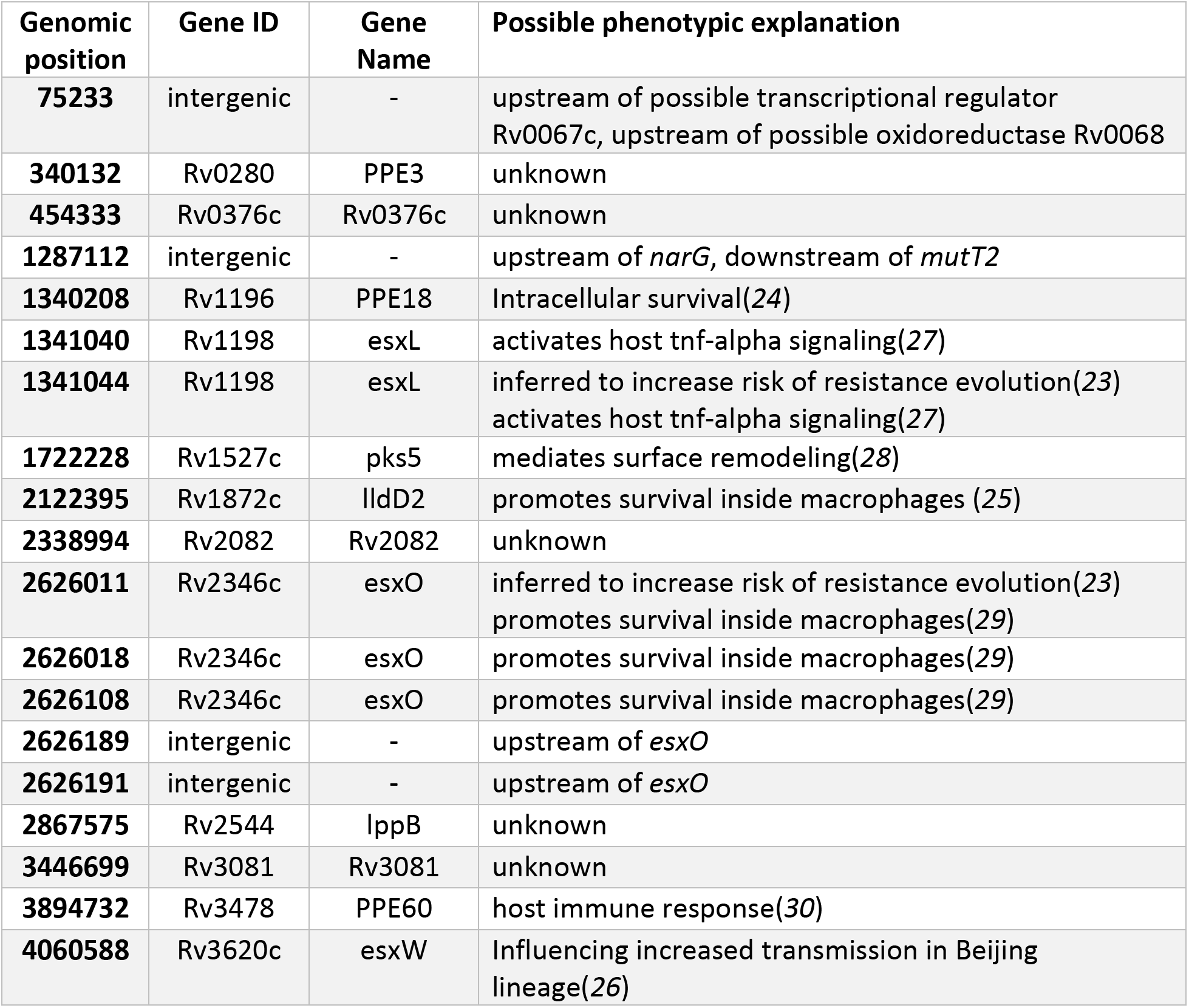
Resistance-potentiating mutations are associated with host-pathogen interactions. Genomic position, identifier, and name for each of 20 mutations found to occur before at least 30 different resistance-conferring mutations. We include a possible phenotypic explanation for the role of each of these potentiating mutations.

We discover several previously implicated SNPs amongst our antibiotic potentiators. This includes position C1341044T (EsxL H13H) and position G2626011A (EsxO I54I, both of which were previously found to increase the risk of resistance evolution (*23*), and four other SNPs in the *esxO* gene body or upstream region. We also identify mutations in proteins known to increase intracellular survival of *M. tuberculosis*, position G1340208A (PPE18 R287Q) and position C2122395T (LldD2 V253M), and a mutation previously associated with increased transmission in the Beijing lineage, position T4060588C (EsxW T2A), to potentiate resistance (*24*, *25*) (*26*).

### Consequential mutations that compensate for or amplify antibiotic resistance

We next focus on dependent mutations occurring as a consequence of the initial evolution of resistance, here called consequential mutations, since these may indicate potential new mechanisms of resistance evolution or compensation for loss of fitness from initial resistance mutations. We detect 1,003 significant dependent mutations after initial resistance mutations, with hits for all 12 antibiotics **(Supplementary Data 5)**. We quantified the prevalence of evolutionarily dependent mutations in resistant isolates in our dataset **(Methods)**. We found that a substantial percentage of strains with initial resistance-causing mutations have sequentially acquired dependencies, ranging from 80% for streptomycin to 9% for fluoroquinolones, indicating a pervasive role in antibiotic resistance evolution. **(Figure 4)**.

**Figure 4:**
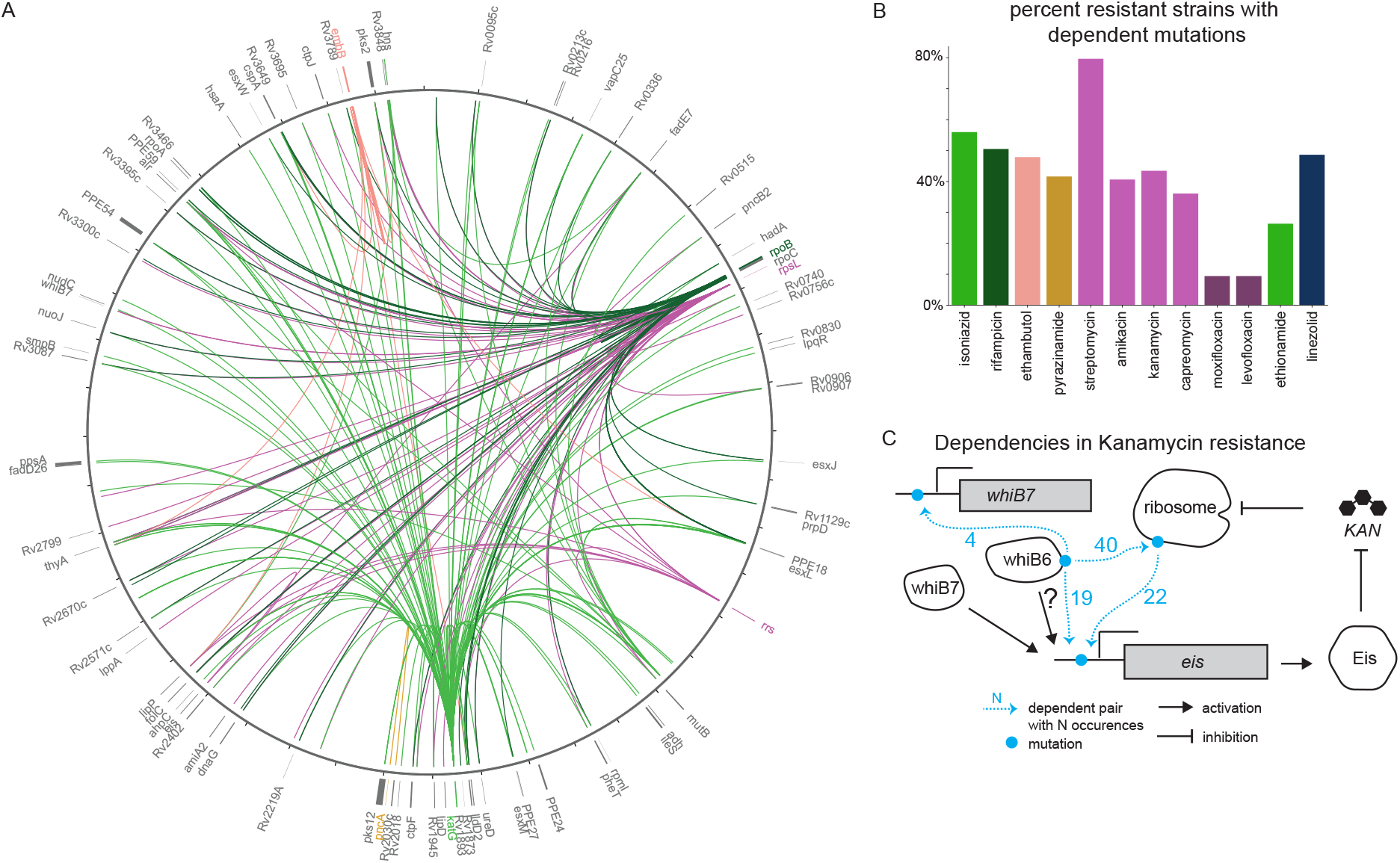
Dependent mutations within resistance-associated genes. We measured the identity and prevalence of significant dependent mutations occurring after initial resistance evolution. (A) Mutations that occur after mutations in known antibiotic resistance genes, visualized on the genome using Circos, with colors corresponding to the antibiotics in panel B. Antibiotics with shared genetic basis of resistance are shown in the same color. Only mutations that happen sequentially at least five times are shown. (B) Fraction of resistant strains that display one or more pairs of sequential dependent mutations. (C) Example of pairs of dependent mutations within the kanamycin resistance pathway, shown on a per-gene basis. Kanamycin’s inhibition of the ribosome is blunted by ribosomal RNA mutations, while cellular kanamycin levels are reduced by increased levels of Eis, putatively caused by both mutations in the *eis* promoter region and mutations in the regulatory proteins WhiB7 and WhiB6. Dependencies between these mutations demonstrate multi-step resistance evolution.

As a positive control, the most frequent consequential mutation we detect is the known strong dependency between rifampicin resistance mutations in RNA polymerase β subunit (RpoB) and substitutions in the RNA polymerase β’ subunit (RpoC), which compensate for the loss of fitness incurred by RpoB mutations through a direct physical interaction (*9*). We also detect dependency between the catalase-peroxidase KatG S315T and position G2726142A, in the *ahpC* gene promoter. Increased levels of the AhpC protein are recognized to compensate for the loss of KatG peroxidase activity (*31*, *32*), demonstrating a possible case of compensatory substitutions mediated by metabolic pathways. The detection of these known relationships reinforces the utility of phylogenetic methods in reconstructing evolutionary dependency.

Our method also detects new relationships. For the aminoglycoside antibiotic kanamycin, we observe consequential mutations likely resulting in amplification of antibiotic resistance between the 16S rRNA gene *rrs*, the target of kanamycin; sites in the promoter region of the N-acetyltransferase Eis, known to degrade kanamycin (*33*, *34*); sites upstream of the transcriptional regulator *whiB7*, known to influence *eis* transcription (*35*); and sites in the transcriptional regulator *whiB6* **(Figure 4)**. Our findings and previous association studies suggest a role for WhiB6 in kanamycin resistance (*36*, *37*). The observed evolutionary dependency suggests that multiple mutations are required to amplify resistance to a high level – mutations in *rrs* disrupt kanamycin binding, while mutations in *whiB6, whiB7*, and *eis* likely increase levels of the Eis protein, leading to increased kanamycin degradation.

Although a gene ontology (GO) analysis did not identify significant enrichment of any one category for non-synonymous mutations following antibiotic resistance **(Methods) (Supplementary Data 6)** (*38*–*40*), one of the top categories is “regulation of DNA-templated transcription”, of major interest since the first-line antibiotic rifampicin targets the RNA polymerase. We find that the RNA polymerase termination factor *nusG* is repeatedly mutated after initial evolution of rifampicin resistance. NusG is notable because it binds directly to the RNA polymerase subunit RpoB (*41*), the target of the drug rifampicin (*42*). The mutated position in NusG, R124H/L, is found at the NusG-RpoB interface **(Methods)**, suggesting that it is involved in stabilizing the action of the mutated polymerase, similar to the compensatory relationship between RpoC and RpoB (*9*) **(Figure 4)**.

A frequent mutation to follow antibiotic resistance in our dataset is HadA C61S, which occurs 40 independent times sequentially or simultaneously with isoniazid resistance evolution and is found in all four major lineages. This mutation is known to confer resistance to the now-obsolete antibiotics thioacetazone and isoxyl (*43*, *44*), and to candidate new antibiotics (*44*, *45*). While the observed HadA mutations are potentially attributable to historical co-administration of thioacetazone and isoniazid (*46*), and hence sequential selective pressure, they may also be consequential mutations of isoniazid resistance -- HadA is upstream of the isoniazid drug target InhA in the mycolic acid biosynthesis pathway (*47*) and may play a role in amplifying isoniazid resistance levels or compensating for InhA mutations.

### Sequential environmental pressures lead to evolutionary dependency in antibiotic resistance

In natural populations, several environmental pressures may act contemporaneously on a population. For pathogenic bacteria, this can take the form of simultaneous or sequential administration of antibiotics to achieve cure. We find strong dependencies between mutations that confer resistance to different antibiotics (**Figure 5**). Notably, this recapitulates the ordering of antibiotic administration in therapy: second-line drug resistance-conferring mutations were consistently acquired on a background of resistance to first-line agents. The observed dependencies also confirm postulated relative fitness costs of resistance mutations for the four first-line drugs (*48*, *49*). These findings demonstrate that evolutionary dependency can be used to study not only molecular dependencies that amplify or stabilize a particular phenotype but also environmental forces when the genetic underpinnings of adaptation are known.

**Figure 5:**
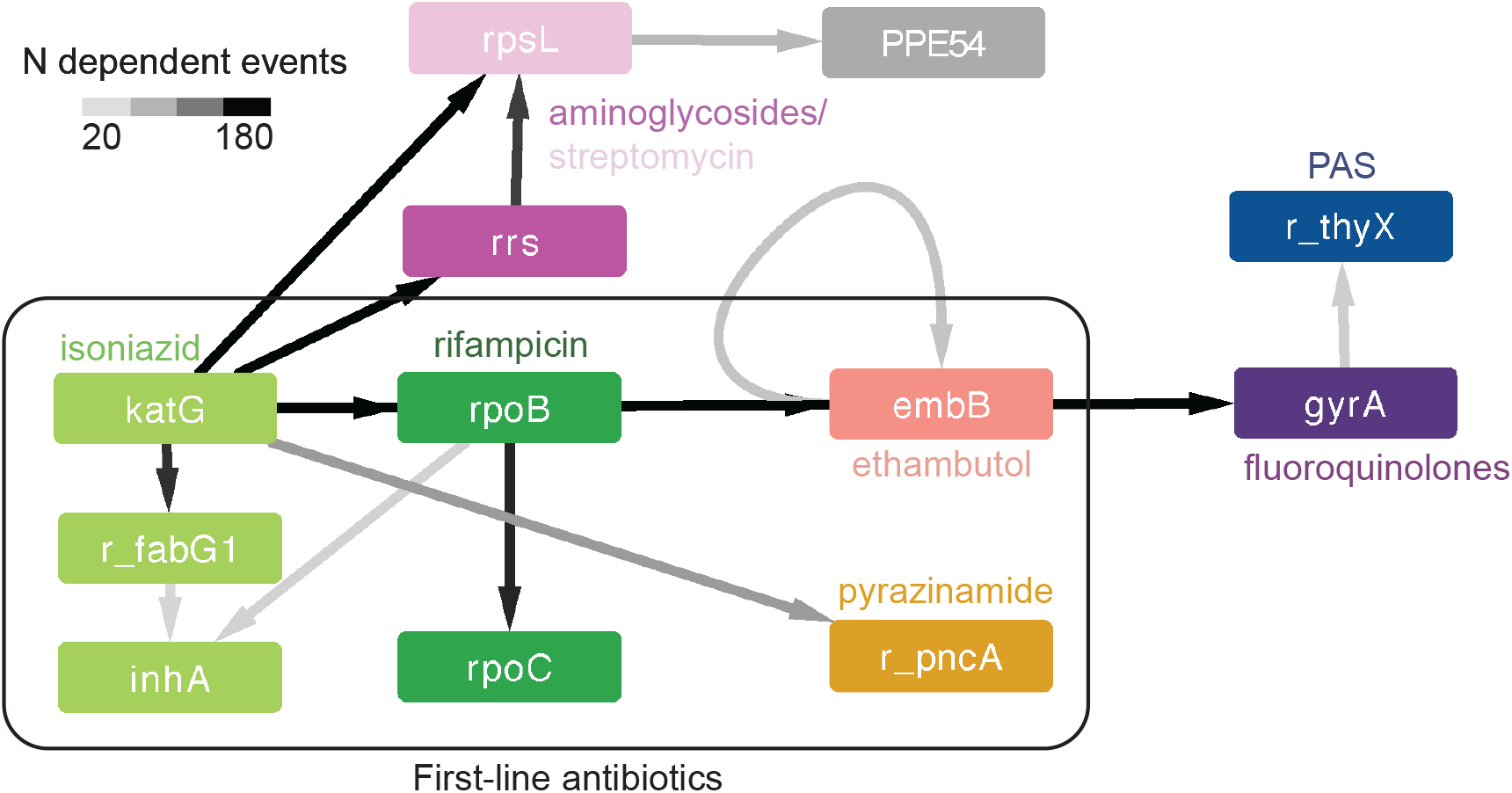
Dependencies between antibiotics. The detected significant dependent mutations between resistance-conferring mutations follow a particular order that mirrors the usage of different antibiotics. For each antibiotic, we took the top dependent pair between known resistance-conferring genes and other genes, and between known resistance genes for different antibiotics. We display pairs and links where mutation *a* occurs sequentially or simultaneously with mutation *b* at least 10 times. The prefix “r_” before a gene name indicates that the mutations are found in the upstream region. The drug para-amino salicyclic acid (PAS) is not included in the WHO catalog, but *thyX* is a candidate resistance gene for this drug(*50*).

### Measuring the effect of dependent mutations on resistance phenotypes

The volume of data available on antibiotic minimum inhibitory concentrations (MIC) limited the statistical power of linear regression for validating direct effects of dependent mutations on antibiotic resistance **(Supplement**). We observe that 95% of the 20 potentiator mutations have a positive, epistatic influence on MIC for at least one drug using this approach, but only a median of 29% of known resistance-conferring mutations were determined to have a statistically significant effect (**Supplement**). We instead examined heritability of MIC and the proportion of variance explained by the dependent mutations using a series of antibiotic specific linear mixed models (with up to n=1,825 observations). MIC is a trait with high heritability – previously estimated at 64%-88% per drug based on all sites in the genome (*37*). Compared to heritability estimated from all homoplastic sites, heritability explained by mutations in known or suspected resistance conferring-genes has a median deficit of 29% per antibiotic **(Table 2)**. Incorporating sites found to have mutation dependencies with antibiotic resistance genes resulted in a median increase of 15% in heritability, accounting for much of the deficit in heritability using 75% fewer sites, despite these dependencies having been derived without phenotypic data. This demonstrates that our proposed mutational dependency analysis is evolutionarily meaningful and characterizes the genetic architecture of antibiotic resistance phenotypes, even if the current analyses lack the power to detect individual sites and pairs as significantly influencing phenotype in a regression analysis.

**Table 2:** Incorporating dependent mutations explains heritability of antibiotic resistance. We compute the heritability of antibiotic MIC using (1) all homoplastic sites in our dataset, (2) homoplastic mutations in known and suspected resistance conferring sites, and (3) homoplastic mutations in known and suspected resistance conferring sites, as well as mutations found to be dependent with known resistance conferring mutations (single sites and interaction terms). Note that “N sites” refers to the number of sites included in the analysis that were actually found to have a polymorphism in the isolates with MIC available.

|  | Homoplastic sites |  | Resistance genes |  | Resistance genes and dependent pairs |  |
| --- | --- | --- | --- | --- | --- | --- |
| Drug | Heritability | N sites | Heritability | N sites | Heritability | N sites |
| Amikacin | 0.70 | 2013 | 0.50 | 102 | 0.54 | 778 |
| Capreomycin | 0.54 | 1665 | 0.28 | 93 | 0.41 | 620 |
| Ethambutol | 0.59 | 2389 | 0.27 | 76 | 0.45 | 885 |
| Ethionamide | 0.55 | 1920 | 0.12 | 61 | 0.47 | 386 |
| Isoniazid | 0.71 | 2393 | 0.22 | 69 | 0.68 | 620 |
| Kanamycin | 0.70 | 1921 | 0.51 | 68 | 0.62 | 744 |
| Moxifloxacin | 0.69 | 1761 | 0.32 | 28 | 0.60 | 50 |
| Pyrazinamide | 0.62 | 1727 | 0.65 | 157 | 0.62 | 371 |
| Rifampicin | 0.66 | 2437 | 0.31 | 152 | 0.59 | 402 |
| Streptomycin | 0.59 | 2397 | 0.33 | 211 | 0.41 | 599 |
| Mean | 0.64 | 1967 | 0.32 | 84.5 | 0.56 | 609.5 |

## Discussion

We propose a new method to uncover evolutionary dependencies between mutations in naturally evolving populations and apply it to 31,435 isolates of *Mycobacterium tuberculosis* complex. We find both sequentially and simultaneously occurring pairs of dependent mutations, which are enriched in antibiotic resistance and antigenic function. We detect 20 potentiating mutations that predispose the evolution of resistance mutations to several antibiotics, and also have a measurable statistical interaction on antibiotic MICs in regression models. We also explore consequential mutations that are acquired in a dependent manner subsequent to resistance acquisition, providing possible examples of novel pathway-mediated selection. We lastly demonstrate the power of this approach in capturing environmental dependencies when the genetic mechanisms are well understood.

We observe that simultaneously occurring mutations are enriched in pairs of mutations within 100 base pairs on the genome occurring in antigenic genes, suggesting that this is due to non-SNP mutational processes such as recombination, which could simultaneously introduce multiple variants in close proximity. Gene conversion has been previously postulated to drive *esx* gene evolution, which are genes enriched in antigenic function (*21*). Innovatively, our results suggest that other genes and especially antigenic genes may evolve through gene conversion, but this requires further validation potentially with long-read sequencing data. After excluding the proximal dependencies, simultaneous distant dependencies are also enriched in antigenic function and in antibiotic resistance. The observation of simultaneous acquisition of antibiotic resistance pairs of variants may relate to the phylogenies’ inability to temporally resolve the two events due to sparse sampling or due to rapid acquisition of the phenotypes in time.

We examine dependent mutations that arise before antibiotic resistance, here called potentiating mutations, or after antibiotic resistance, here called consequential mutations. Consequential mutations appear to fall into at least two categories, those that compensate for loss of fitness due to resistance acquisition, and those that amplify the phenotype of antibiotic resistance itself. For example, *nusG* mutations appear to compensate for destabilizing *rpoB* mutants based on our structure analysis, and *hadA* knockdowns were found to significantly sensitize strains to high levels of isoniazid in a recent CRISPRi study (though the magnitude of depletion was below study threshold) (*51*). In contrast, the mechanism by which our 20 observed potentiating mutations predispose the evolution of antibiotic resistance is still in question. One possibility is that proteins on the cell surface, including antigenic proteins, play a direct role in antibiotic resistance, for example through altering cell permeability. This possibility is supported by the observation that 95% of observed potentiating mutations have a detectable epistatic influence on MIC. Another possibility is that strains with potentiating mutations may be more likely to transmit between hosts or progress from latent to active tuberculosis disease, leading to higher exposure to antibiotic treatment. Strains with potentiating mutations may also reach higher effective population sizes within host, leading to higher probability of resistance evolution. Finally, strains with potentiating mutations may have higher overall fitness, preemptively compensating for loss of fitness due to resistance evolution.

We investigated whether the detected dependencies were associated with higher antibiotic resistance levels as measured by strain minimum inhibitory concentrations (MICs). Dependent mutations when added to known resistance-conferring variants capture the majority of heritability, and several mutations including 95% of the 20 potentiating mutations have measurable associations on resistance. As more MIC data becomes available, we expect that the power of these analyses to capture the individual effects of dependent mutations will improve.

Our method is not without limitations. It relies on repeated observations of evolutionary events to infer significant non-independence of mutations. Therefore, its power is dependent on the number of times a mutation has arisen, and thus is biased against the effects of very recent selection, e.g. responses to newer antibiotics, such as linezolid, clofazimine, and even fluoroquinolones. The smaller numbers of dependent mutations observed for these drugs should not be taken as an assertion that there are fewer dependent mutations as a result of the evolution of resistance to these drugs, but rather that we do not yet have enough observations of evolutionary trajectories to reliably infer significance. This issue is also present in the case of pyrazinamide, where a large number of variants in the *pncA* gene are known to cause resistance, and thus the statistical signal is diluted over a large number of variants. A future extension to address this limitation is the expansion to study dependence between mutational burden measured per gene or regions. Lastly, the links inferred by our method are based only on the presence of pairs of mutations, and thus captures associations due to biological epistasis or other forces like sequential or simultaneous environmental pressures.

We believe the method introduced here will be readily generalizable to other microbial species. While *M. tuberculosis* generally does not participate in horizontal gene transfer and thus our method focused on SNPs, our framework could extend to analyzing not just the probability of individual mutations but the probability of gene acquisition or other mutation events. Our method has broad conceptual applicability to understanding clonal evolution ranging from viruses to cancer cells. We show that in *M. tuberculosis*, dependent mutational events are enriched in mutations associated with antibiotic resistance and antigenic function. We discover 20 mutational events that appear to potentiate antibiotic resistance, and dependent events arising as a consequence of resistance are due to both compensatory variation and amplification of resistance phenotypes. Together these results represent a wealth of new knowledge about the evolution of an important microbial pathogen.

## Methods

### Dataset of *M. tuberculosis* strains

We use a previously curated dataset of 31,428 strains, representing six major *M. tuberculosis* lineages (*16*). Whole genome sequence data is processed using a previously validated pipeline (*52*, *53*) Briefly, reads are aligned to the H37Rv reference genome using BWA-MEM after trimming and filtering with PRINSEQ and contaminant removal with Kraken (*54*–*56*). Variant calling is performed with Pilon and duplicate reads were removed using Picard (*57*, *58*). All isolates had at least 95% coverage of the reference genome at 10x coverage. We excluded regions of H37Rv with mappability <0.90 (*59*).

### Reconstructing mutational history

We use a dataset that reconstructed the evolutionary history of the 31,428 strains by building maximum likelihood phylogenies and performing maximum likelihood ancestral reconstruction (*60*) to infer mutational events over time in the population of strains (*16*). We consider only single nucleotide substitutions (SNS) in the coding and non-coding portions of the genome. We annotate each single nucleotide polymorphism as either *to* or *from* the ancestral state based on an inferred ancestor of extant the *M. tuberculosis* complex (*17*).

To study mutational dependencies, we consider non-ancestral pairs of mutations that occurred either sequentially or simultaneously. Simultaneous mutations are pairs inferred to have occurred on the same branch of the tree (note that these mutations may not have actually occurred simultaneously in a single mutation event, but their ordering cannot be resolved). Sequential mutations are pairs are inferred to have occurred on different, sequential branches.

### A model to detect evolutionary dependency between sequentially occurring mutation pairs

We model the probability of a given non-ancestral mutation, ***a***, in the presence or absence of a second mutation, ***b***, as follows: In the phylogenetic tree with N branches, we define the Bernoulli random variable X to indicate whether a mutation occurs on a particular branch. For example, *X_a,n_* = *1* if ***a*** evolves on the ***n***th branch and *X_a,n_= 0* if mutation ***a*** does not occur on the ***n***th branch. We define the Bernoulli random variable Y to indicate whether a mutation has already evolved prior to a particular branch. For example, *Y_b,n_ = 1* if ***b*** evolved prior to the ***n***th branch, and *Y_b, n_= 0* if ***b*** did not evolve prior to the ***n***th branch.

We model the probability of ***a*,** P(X_a,n_), as a Beta distribution, the conjugate prior of the Bernoulli distribution. The shape parameters α and β of the Beta distribution are given by the count of observed branches where X_a,n_=1 and X_a,n_=0, respectively,

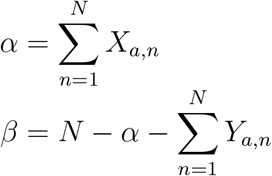

Branches where mutation *a* has already occurred (*ie*, Y_a,n_ = 1) are subtracted because there is no possibility of further mutation in our model. Branch length is not considered in this calculation.

Because we seek to test whether P(X_a_|Y_b_=1) is different from P(X_a_|Y_b_=0), we partition the branches into two sets: those with mutation ***b***, {L | Y_b,l_=1 or X_b,n_=1}, and those without mutation ***b*** {M | Y_b,m_=0 and X_b,n_=0}. To test whether the two distributions are sufficiently different, we test the hypothesis that the expected value of Beta(α_M_, β_M_) is drawn from Beta(α_L_, β_L_) by computing the p-value. We include a pseudocount of 1 for all values of alpha and beta. This approach to modeling P(X_a_) using the observed mutation data captures the higher uncertainty about P(X_a_|Y_b_=1) when the number of branches in {L} is small, because the variance of the beta distribution is higher for smaller values of α and β.

### Detecting evolutionary dependency between simultaneously occurring mutations

To determine whether two mutations ***a*** and ***b*** occur simultaneously more often than expected, we again model the probability of their simultaneous occurrence using a beta distribution where:

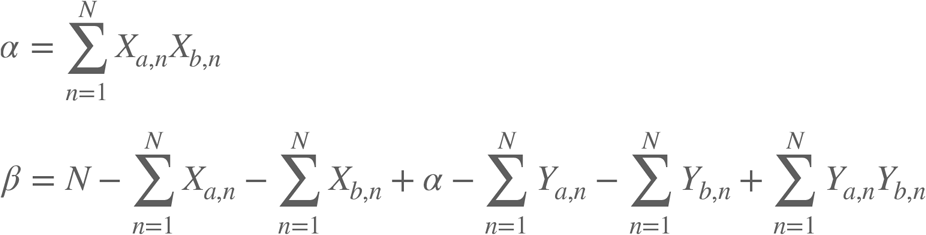

Alpha is the number of branches where both mutations occur, and beta is the number of branches where neither mutation occurs and neither mutation has already occurred.

The null expectation for the frequency with which mutations occur on the same branch is based on the individual frequency of the mutations:

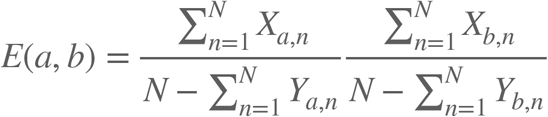

We then determine the probability of drawing the null expectation from the estimated distribution of co-occurrence probability, which constitutes the p-value.

### Assigning mutations to functional categories

We define a set of antibiotic resistance-associated sites based on World Health Organization data for 12 antituberculosis antibiotics (*18*) **(Supplementary Data 4)**. We also include the entire intergenic regions upstream and downstream of each resistance-associated gene, using gene location data from Mycobrowser (*61*), because non-coding regions can have substantial effects on resistance phenotypes(*37*).

We define a set of antigenic genes based on data from IEDB (*20*). Following previous work on *M. tuberculosis* antigens (*19*), a list of all antigens was downloaded on January 7^th^, 2022 with the following query criteria: linear peptide, Organism= “mycobacterium tuberculosis complex” (ID 77643), positive assays only, T-cell binding, any MHC restriction class, human host, any disease type, any reference type. Any gene present in this list is considered antigenic.

To define gene essentiality, we take the union of all essential genes listed in Table S3 of Minato et al, 2019, which summarizes the results of three studies on gene essentiality (*62*).

We use two previous studies to define positions that contain lineage-associated variants (*53*, *63*).

### Defining resistance-potentiating mutations

We define any sequential mutation pair where the second mutation is a known resistance-conferring mutation and the first mutation is not as resistance-potentiating. After observing that certain initial mutations occur before many different resistance-conferring mutations, we focus our analysis on these “most potentiating” initial mutations by selecting only those where the initial mutation is followed by over 30 different resistance-conferring mutations.

### GO terms

We performed Gene Ontology Enrichment analysis against the *Mycobacterium tuberculosis* genome using the Gene Ontology website (geneontology.org, release date 2021-05-01) (*38*–*40*).

### Computing heritability

Heritability calculations were run using GEMMA v0.98.1 (*64*). For antibiotics tested in media other than 7h10, MIC values were normalized by dividing by the ratio of the critical concentration in 7h10 to the critical concentration in the tested media. MIC values were converted from a range to a number by taking the midpoint of the range, or the endpoint if only one point was provided (eg, “>10” becomes “10”, “2-4” becomes “3”), and then were log-transformed. Alleles were encoded as 0 for ancestral state, 1 for non-ancestral, or missing for positions where the allele could not be confidently called. For each set of sites, the sites of interest were used to define a GRM, and the proportion of variance (PVE) explained by the GRM was calculated. This is equivalent to the heritability.

Three sets of sites were tested: (1) All homoplastic sites (sites with at least 5 independent mutations), (2) Homoplastic mutations in a known or suspected resistance-conferring site, and (3) Homoplastic mutations in a known or suspected resistance-conferring site, or found to be a dependent mutation with a mutation in a known resistance-conferring site (including both single and pair terms).

### Determining physical distance between nusG and rpoB mutations

We downloaded the solved structure of the RNA polymerase – NusG complex (PDB ID: 6z9p) (*41*) from the RCSB PDB (*65*). We used MUSCLE from the EBI webserver with default parameters (*66*)to align the sequence of *M. tuberculosis* NusG and RpoB to the sequence of the crystal structure. Then, we used PyMOL v2.4.0 to measure the physical distance between the residue corresponding to *M. tuberculosis* position 734624 (NusG R124), to any residue in the RpoB protein (*67*).

## Supporting information

Supplementary Data 1 - Homoplastic SNPs

Supplementary Data 2 - Chi-Squared Tests

Supplementary Data 3 - Significant Dependent Pairs

Supplementary Data 4 - Resistance-conferring mutations

Supplementary Data 5 - Dependent mutations implicated in resistance

Supplementary Data 6 - GO enrichment

## Acknowledgements

We thank the members of the Farhat lab for their discussions and insight. We thank Dr. Lisa Lojek and Dr. Chris Sassetti for initial attempts to construct *hadA* mutant strains. Computational resources and support were provided by the Orchestra High Performance Compute Cluster at Harvard Medical School, which is funded by the NIH (NCRR 1S10RR028832-01). AGG was supported by a National Institutes of Health NLM Training Grant T15LM007092 and NIH/NIAID F32AI161793. R.V.J. was supported by the National Science Foundation Graduate Research Fellowship under Grant No. DGE1745303.

