## Supplement for "The discovery of genome-wide mutational dependence in naturally evolving populations"

**Potts model results dominated by lineage variants and non-SNP mutational processes in *M. tuberculosis***

While Potts models have proven utility for detecting epistatic pairs in *Streptococcus pneumoniae, Streptococcus pyogenes, and Neisseria gonorrhoeae* (*1–3*), we find that in the particular case of *M. tuberculosis*, the results are dominated by putative non-SNP mutational processes and lineage-specific variants. Here, we define a non-SNP mutational process as any mutational event that generates more than one single nucleotide polymorphism at a time, either by substituting multiple bases or through recombination, which can manifest as apparent multiple substitutions after read mapping.

We initially ran a Potts model on all non-synonymous mutations with an allele frequency greater than 0.001, using the SuperDCA software package (*4*). To reduce the dataset size, we used only 11,015 isolates from our larger set of 31,435. We called variants relative to the reference genome of H37Rv, where any sites that show an alternative base in more than 40% of reads were called as polymorphisms. We translated the genome and selected only sites with non-synonymous variants, for a total of 36,489 sites. SuperDCA, was run using a minor allele frequency threshold of 0.001 (for a total of 10,278 sites) and no-reweighting, with all other parameters set to default values. Following the SuperDCA workflow, we re-ranked the couplings based on phylogenetic weighting using HierBAPS(*2*, *5*). Top-ranking couplings were chosen by fitting a linear model where the couplings were used to predict the log10 of coupling rank, and selecting points with a residual greater than 5 times the estimated standard deviation, for a total of 201,789 significant pairs out of 42,980,356 (*1*, *2*) (**Supplementary Figure 1**). Despite following the literature standard procedure for phylogenetic weighting and determining significant hits, we found that the majority of the top-scoring hits were lineage-associated, and a substantial fraction were found in the same gene, potentially indicating evolution due to a single mutational event **(Supplementary Figure 2, Supplementary Table 8)**.

To better control for population structure, we then ran a Potts model on just the homoplastic sites in our analysis. Potts models were run on our set of 4,776 homoplastic variants from 31,435 isolates using the plmc package (*3*, *6*) with a maximum of 200 iterations. Alleles were encoded with three states: ancestral, derived, or gap (used for both deletions relative to the reference and uncertain allele calls). No sites in the alignment had more than 10% gaps. We scanned a range of theta values (0.01, 0.02, 0.05, 0.08, and 0.1) to determine a value which sufficiently corrects for oversampling of certain lineages without overcorrecting **(Supplementary Table 9)**. We chose theta = 0.02, which produces a N_eff of 3623.7. As in Schubert and Maddamsetti *et al*, we used a two-component mixture model to select the strongly coupled pairs, finding 33,647 pairs (out of a total of 11,255,140) over the 99% probability threshold (*3*, *7*). Couplings were processed using the EVCouplings Python package (*8*). To further control for population structure, we removed any pairs with a member in a lineage-associated position (*9*, *10*), leaving 28,374 sites.

We find that while this protocol did correct for lineage-associated mutations, the model still has results dominated by mutations due to a single mutational event (**Supplementary Figure 3**). The top pairs tended to be in close genetic proximity – 90% of the top 500 hits are within 100 base pairs on the genome. While this may be explained by true evolutionary dependencies due to shared function of proximal base pairs, it may also be explained by multi-base mutations, gene conversion, or recombination events. The ancestral sequence reconstruction analysis supported the latter possibilities: 54% of the top 500 pairs are predicted to arise on the same phylogenetic branch more than 80% of the time and were not sequential events, and a further 16% are found in positions with more than 5% gaps, indicating possible insertions and deletions (**Supplementary Table 10**). The enrichment for non-SNP evolutionary events is potentially due to the focus of Potts models on isolate genomes: mutations that affect multiple sites manifest as multiple SNPs that always co-occur and are never found independently, generating strong signal.

**Linear mixed models for detecting direct effects of dependent mutations on antibiotic minimum inhibitory concentrations**

We measure which dependent mutations have a direct effect on antibiotic resistance, by running a series of linear mixed models of antibiotic minimum inhibitory concentration (MIC), including linear (additive) and interaction (epistatic) terms of each pair of variants (2,371 sites and pairs tested on data from n=1,825 isolates).

Association tests were run using GEMMA v0.98.1 using LMM mode and a missing allele threshold of 20% (*57*). Minimum inhibitory concentration (MIC) data was obtained by combining data from multiple studies spanning 2124 isolates (*30*, *45*, *58*–*60*). For antibiotics tested in media other than 7h10, MIC values were normalized by dividing by the ratio of the critical concentration in 7h10 to the critical concentration in the tested media. MIC values were converted from a range to a number by taking the midpoint of the range, or the endpoint if only one point was provided (eg, “>10” becomes “10”, “2-4” becomes “3”), and then were log-transformed. Alleles were encoded as 0 for ancestral state, 1 for non-ancestral, or missing for positions where the allele could not be confidently called. Each evolutionarily dependent pair of alleles was tested in a single multivariate linear mixed model, which included an interaction term to capture epistatic effects. We controlled for population structure using a GRM computed using all alleles (not just homoplastic variants) with a minor allele frequency greater than 0.1% across all 31,435 isolates in our dataset.

The percent of dependent events with a detectable influence on MIC, either additively or epistatically, ranged from 3% for pyrazinamide to 40% for moxifloxacin, with a median of 12%. **(Supplementary Table 1).** Notable examples include a promoter variant in position 4243217 in the *embCAB* locus with a positive linear influence on ethambutol MIC, and a synonymous variant in position 332951 (VapC25 P62P) with a measure positive epistatic influence on rifampicin and isoniazid resistance **(Supplementary Table 2, Supplementary Data 7)**. VapC25 is a toxin suggested to promote antibiotic tolerance by slowing growth rate in host (*43*). We observe that 95% of the 20 potentiator mutations have a positive, epistatic influence on MIC for at least one drug. We expect that as more phenotypes become available, more variants will have detectable influences on MIC – currently only a median of 29% of known resistance-conferring mutations were determined to have a detectable statistical influence on resistance, indicating that greater power is needed to detect all effects.

**Supplementary Figure 1: Using a semi-log linear model to determine significant couplings from SuperDCA**

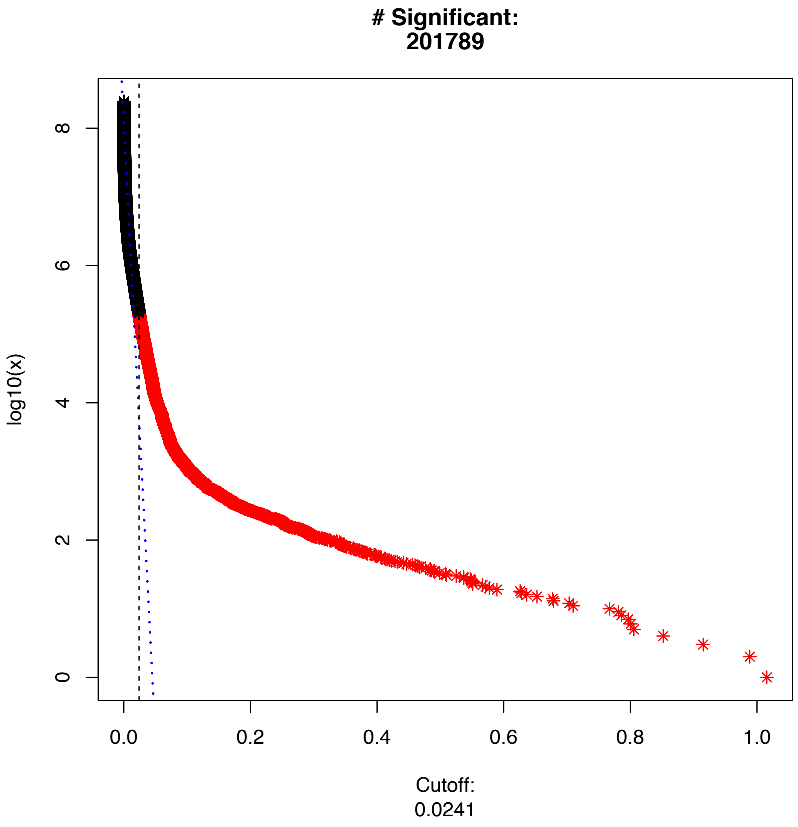

**Supplementary Figure 2: The top-scoring hits from SuperDCA are lineage-associated or single mutational event changes.** The top 10,000 ranked hits from SuperDCA, colored by their category, are shown. Mutation pairs are defined as in_lineage if at least one mutation is lineage-associated according to HierBAPS. Mutation pairs that are not in_lineage or found in the same gene are considered "standard". Only the top 10,000 hits out of 201,789 are shown for visualization purposes.

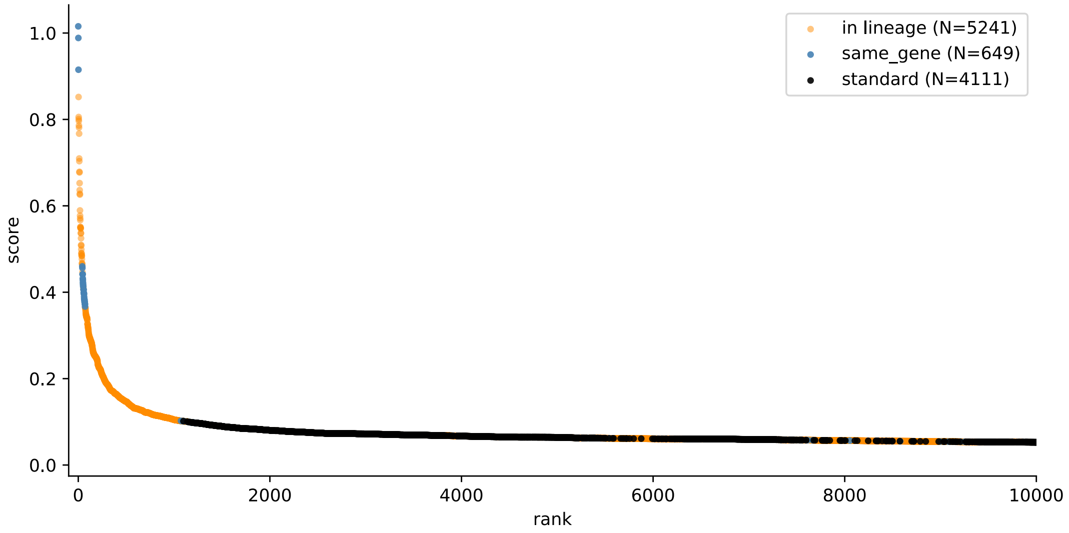

**Supplementary Figure 3: Identity of top 500 Potts model hits among homoplastic sites.** The identity of the top-ranking pairs of couplings output by the Potts model. Sites are labelled as "standard" if they are not inferred to be part of the same mutational event and neither of the sites have >5% gaps in the input sequence alignment.

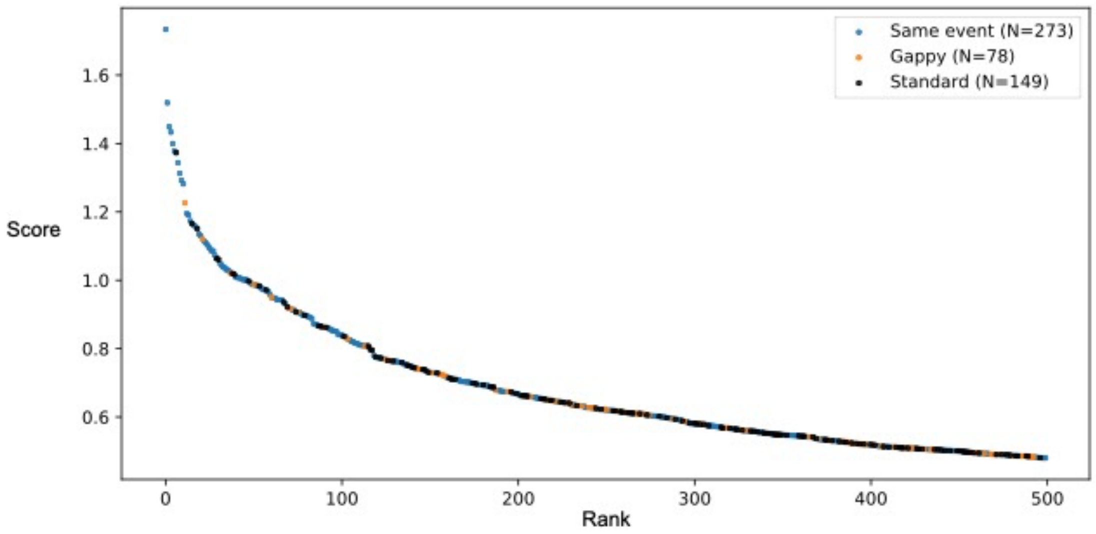

**Supplementary Table 1: Percent of known and dependent mutations with significant effect on MIC**

| | Known resistance variants | | | Dependent mutations | | |
|---|---|---|---|---|---|---|
| drug | Number tested | Number significant | Percent significant | Number tested | Number significant | Percent Significant |
| AMIKACIN | 10 | 1 | 10% | 159 | 24 | 15% |
| CAPREOMYCIN | 7 | 3 | 43% | 146 | 20 | 14% |
| ETHAMBUTOL | 11 | 4 | 36% | 182 | 19 | 10% |
| ETHIONAMIDE | 20 | 3 | 15% | 83 | 7 | 8% |
| ISONIAZID | 7 | 3 | 43% | 178 | 7 | 4% |
| KANAMYCIN | 10 | 3 | 30% | 152 | 25 | 16% |
| MOXIFLOXACIN | 9 | 6 | 67% | 98 | 37 | 38% |
| PYRAZINAMIDE | 86 | 5 | 6% | 174 | 6 | 3% |
| RIFAMPICIN | 21 | 6 | 29% | 256 | 17 | 7% |
| STREPTOMYCIN | 27 | 4 | 15% | 260 | 35 | 13% |

**Supplementary Table 2: Consequential mutations with significant effect on MIC**

Mutations occurring after antibiotic resistance (consequential mutations) that have a significant influence on MIC, either linearly or epistatically. Variants and genomic positions indicated with a '+' are epistatic interactions between the two variants that are found to be significant. Genes indicated as X-Y are found in the intergenic region between genes X and Y.

| Gene(s) | Genomic position(s) | drug | beta | p_wald |
|---|---|---|---|---|
| hadA | 1473246 | KANAMYCIN | 1.161 | 0.008 |
| hadA + rrs | 1473246 + 732110 | KANAMYCIN | 1.748 | 0.009 |
| inhA + PPE19 | 1674481 + 1532777 | ETHIONAMIDE | 0.987 | 0.002 |
| katG + vapC25 | 2155168 + 332951 | ISONIAZID | 1.421 | 0.001 |
| katG + PE_PGRS28–Rv1453 | 2155168 + 1638364 | ISONIAZID | 1.082 | 0 |
| katG + Rv1873 | 2155168 + 2123182 | ISONIAZID | 1.372 | 0 |
| rpoB + vapC25 | 761110 + 332951 | RIFAMPICIN | 1.577 | 0.001 |
| embB + PE_PGRS28–Rv1453 | 4247429 + 1638364 | ETHAMBUTOL | 0.297 | 0.002 |
| embC–embA | 4243217 | ETHAMBUTOL | 0.391 | 0.005 |
